# Estradiol regulates voltage-gated potassium currents in corticotropin-releasing hormone neurons

**DOI:** 10.1101/2023.01.16.524323

**Authors:** Emmet M. Power, Dharshini Ganeshan, Karl J. Iremonger

## Abstract

Corticotropin-releasing hormone (CRH) neurons are the primary neural population controlling the hypothalamic–pituitary–adrenal (HPA) axis and the secretion of adrenal stress hormones. Previous work has demonstrated that stress hormone secretion can be regulated by circulating levels of estradiol. However, the effect of estradiol on CRH neuron excitability is less clear. Here we show that chronic estradiol replacement following ovariectomy increases two types of potassium channel currents in CRH neurons; fast inactivating voltage-gated A-type K^+^ channel (I_A_) currents and non-inactivating M-type K^+^ currents (I_M_). Despite the increase in K^+^ currents following estradiol replacement, there was no overall change in CRH neuron spiking excitability assessed with either frequency-current curves or current ramps. Together, these data reveal a complex picture whereby ovariectomy and estradiol replacement differentially modulate distinct aspects of CRH neuron and HPA axis function.

**Summary statement:** Chronic estradiol replacement in ovariectomised mice influences voltage-gated potassium channel function.

## Introduction

Corticotropin-releasing hormone (CRH) neurons in the paraventricular nucleus (PVN) of the hypothalamus are neuroendocrine neurons which control activity of the hypothalamic-pituitary-adrenal (HPA) axis ^1–7^. These neurons are activated in response to stress ^3^ which leads to CRH secretion from the median eminence into the portal circulation. This triggers secretion of adrenocorticotropic hormone (ACTH) from the anterior pituitary which subsequently stimulates corticosteroid synthesis and release from the adrenal cortex.

Activity of the HPA axis is sexually dimorphic. In rodents, females have higher levels of circulating corticosterone as well as stress-evoked corticosterone release ^8^. There are also marked changes in activity of the HPA axis across the female reproductive cycle with both basal and stress-evoked levels of corticosterone being highest on proestrus ^9,10^. Since estradiol is a primary sex hormone in females and is at its highest levels on proestrus, this has led to the theory that estradiol could be responsible for both sex and estrous cycle differences in HPA axis activity ^11^. Consistent with this idea, basal corticosterone secretion in female rats is reduced following ovariectomy ^8,12,13^. Circulating corticosterone levels can also be elevated in ovariectomized (Ovx) rats with subsequent estradiol replacement ^14–16^. Despite this, other data in rats show that estradiol suppresses stress-evoked ACTH release ^12,13^ as well as stress-evoked cFos labelling in CRH neurons ^14,17,18^. To add to this complex picture, studies investigating the effect of estradiol replacement in Ovx mice are conflicting. Some studies report that estradiol replacement in Ovx mice can increase corticosterone levels ^19^, while others report no effect ^20–22^ or even reduced corticosterone ^23–27^. Overall, the impact of estradiol on HPA axis function and CRH neuron activity is complex and may differ between species.

We have recently shown that K^+^ channel function and CRH neuron excitability are regulated over the estrous cycle in mice ^28^. During the proestrus phase of the estrous cycle, coinciding with a peak in estradiol levels, CRH neurons exhibit smaller K^+^ channel currents and higher levels of excitability measured by electrophysiological recordings. This corresponds with previous publications showing basal and stress-evoked corticosterone levels being highest during proestrus ^9,10^. However, it is currently unclear if these changes in excitability are driven by estradiol alone. Previous work has shown that estradiol can regulate K^+^ currents and excitability in other central neurons ^29–31^. Therefore, in this current study, we aimed to determine the effect of Ovx and high or low levels of estradiol replacement on K^+^channel function and CRH neural excitability. Using patch-clamp recordings from CRH neurons, we show that chronic elevations in estradiol levels in Ovx female mice leads to increased levels of two K^+^channel currents: I_A_ and I_M_. However, chronic estradiol elevations did not significantly change intrinsic excitability compared to Ovx animals. This suggests that the magnitude of changes in K^+^ channel currents were not sufficient to impact spiking excitability. Despite this, we speculate that enhanced K^+^channel function may impact how these neurons integrate and process stress relevant synaptic inputs.

## Methods

### Animals

All electrophysiological experiments were carried out in adult female (2-6 months old) Crh-IRES-Cre;Ai14 (tdTomato) mice. These mice were generated by crossing Crh-IRES-Cre (B6(CG)-Crhtm1(cre)Zjh/J)^32^ strain with Ai14(B6.Cg-Gt(ROSA)26Sortm14(CAG-tdTomto)Hze/J) strain, both originally obtained from Jackson laboratories (stock numbers 012704 and 007914, respectively). These mice have been previously shown to faithfully label CRH neurons in the PVN ^33–35^. Serum corticosterone and tissue samples were taken from a mixture of C57/Bl6J (Jackson laboratory) and Crh-IRES-Cre mice (2-4 months). Animals had a 12h light/dark cycle (7am-7pm lights on) with food and water available *ad libitum*. All protocols and procedures were approved by the University of Otago Animal Ethics Committee and carried out in accordance with the New Zealand Animal Welfare Act.

### Ovariectomy and hormone replacement

Adult female mice (> 3 months) were bilaterally ovariectomized under isoflurane general anesthetic. Simultaneously mice received a 10 mm long silastic capsule (inner diameter: 1.57 mm; outer diameter: 2.41 mm) containing 17β-estradiol (estradiol) subcutaneously implanted between the shoulder blades and neck. The dose of estradiol (E8875, Sigma) was based on previous publications and estimated to give levels similar to estrus/diestrus for the Ovx^LowE^ group and proestrus (or higher) for the Ovx^HighE^ group ^36–38^. Ovx^LowE^ mice received an implant with 4 μg estradiol dissolved in absolute ethyl alcohol and mixed with silastic gel. Ovx^HighE^ mice received an implant containing crystalline estradiol mixed 1:1 with cholesterol. One group of mice were Ovx and received an implant containing only cholesterol. All mice were left for 2-3 weeks before being used for tissue collection or electrophysiology.

### Blood, tissue collection and ELISA

All mice were habituated to handling for at least 4 days prior to tissue collection. Mice were euthanised (between 9-11am) and trunk blood was collected in tubes. All blood samples were kept on ice before being centrifuged. Uterus and adrenal glands were dissected out and weighed immediately following decapitation. Uterus weights were also taken from a subset of animals used for electrophysiology, protocol for dissection and weighing remained the same. Adrenal gland weight is the combined weight of both left and right adrenals for each animal. Thymus glands were dissected out and stored in 4% PFA before being weighed. Serum corticosterone was measured using an ELISA (Arbor Assays, Cat# K014,RRID AB_2877626) according to the manufacturer’s instructions.

### Slice preparation

Mice were killed by cervical dislocation between 9-11 am, their brain quickly removed and placed in ice-cold oxygenated (95% O_2_, 5% CO_2_) slicing solution containing (in mM); 87 NaCl, 2.5 KCl, 25 NaHCO_3_, 1.25 NaH_2_PO_4_, 0.5 CaCl_2_, 6 MgCl_2_, 25 D-Glucose, 75 sucrose, pH 7.2-7.4. A vibratome (VT1200S, Lecia Microsystems) was used to cut 200 μm-thick coronal slices of the PVN, which were then incubated in oxygenated artificial cerebrospinal fluid (aCSF) containing in (mM); 126 NaCl, 2.5 KCl, 26 NaHCO_3_, 1.25 NaH_2_PO_4_, 2.5 CaCl_2_, 1.5 MgCl_2_, 10 D-Glucose at 30°C for at least 1 hour before recording. For recording, slices were transferred to a recording chamber and continuously perfused with 30°C aCSF at 1.5ml min^-1^. CRH neurons within the PVN were visualized using a 40x objective and epifluorescence to excite tdTomato.

### Whole-cell electrophysiology recordings

Electrophysiological recordings were collected with a Multiclamp 700B amplifier (Molecular Devices), filtered at 2 kHz, and digitized using the Digidata 1440a (Molecular Devices). Data were analysed with Clampfit 10.7 (Molecular Devices).

For whole-cell recordings, borosilicate glass pipettes (tip resistance: 2-5 MΩ) were filled with an internal solution containing (in mM): 120 K-gluconate, 15 KCl, 0.5 Na_2_EGTA, 2 Mg_2_ATP, 0.4 Na_2_GTP, 10 HEPES, 5 Na_2_-phosphocreatine and 0.25% Neurobiotin (adjusted to pH 7.2 with KOH; adjusted to ≈290 mOsm with sucrose). All current clamp experiments were performed in the presence of 10 μM cyanquixaline (6-cyano-7-nitroquinoxaline-2,3-dione) (CNQX) and picrotoxin (50 μM). Each cell was held at approximately −60 mV. The liquid junction potential was calculated to be approximately −14.1 mV and was not compensated for. Cells were not recorded from if input resistance was below 0.7 GΩ or access resistance was above 30 MΩ and both input and access resistance were monitored throughout to ensure stable recording. We used a current step protocol to determine spike output and first spike latency (FSL). The step protocol consisted of 300 ms square steps from 0 to +50 pA in 5 pA increments. Spikes were detected using a threshold search in Clampfit and were analysed for rise time, decay time, amplitude and half width. FSL was calculated from the time of the depolarizing step initiation to the action potential (AP) threshold for the first spike evoked at steps equal or greater than 10 pA. AP threshold was defined as the voltage at which the AP first derivative crossed 10 mV/ms. The same analysis criteria were used to identify FSL and AP threshold for a 1 sec, +40 pA/s ramp protocol.

For all voltage clamp recordings neurons were clamped at −60mV. Input resistance, access resistance and capacitance were monitored periodically throughout recordings. I_A_ current recordings were performed in the presence of CNQX (10 μM), picrotoxin (50 μM), Tetrodotoxin (TTX) (0.5 μM), XE991, 40 μM and nifedipine (100 μM). To evoke I_A_ currents, neurons were hyperpolarized from −60 mV to −110 mV for 500 ms before a family of depolarizing steps were delivered in 10 mV steps from −100 mV to +30 mV. Peak I_A_ amplitude for each voltage step was measured and normalized to capacitance to give the current densities (pA/pF).

A protocol was used to measure the I_M_ relaxation current, similar to that used in previous studies ^39,40^. These recordings were performed in the presence of CNQX (10 μM), picrotoxin (50 μM) and TTX (0.5 μM). This protocol consisted of a pre-pulse to −20 mV for 300 ms followed by 500 ms steps from −30 to −75 mV. The I_M_ relaxation current was measured as the amplitude difference between the initial current and the sustained current at the end of the voltage step.

### Analysis

Statistical analysis was performed using GraphPad Prism 8. All reported values are the mean ± SEM. Comparisons between groups were carried out using either One or Two-way ANOVA where appropriate, with Tukeys post hoc multiple comparison tests. All n-values represent neuron number, all groups had N > 3 animals. *P*<0.05 was considered statistically significant. P values reported on figures are for post hoc multiple comparison tests.

## Results

### Increase in uterine weight but no change in adrenal weight or basal morning corticosterone level following chronic estradiol manipulation

In order to manipulate estradiol levels, female mice were ovariectomized and either received no treatment (Ovx) or received a low (Ovx^LowE^) or high (Ovx^HighE^) dose estradiol implant. Two to three weeks later, animals were euthanized and blood and tissue were collected. The uterus is highly sensitive to estradiol, shows enlargement in response to estradiol elevations and has been previously used as a bioassay for estrogen levels ^41,42^. Estradiol treatment induced a significant increase in uterine weight (One-way ANOVA, F_(2,32)_ = 51.91, *P* < 0.0001; Fig 1A) consistent with previous studies. Previous work has shown that estradiol treatment can also elevate basal corticosterone levels in rats ^14,16,43^ but have no effect ^20–22^ or reduce corticosterone levels in mice ^23–25^. Here we find that in Ovx mice, high or low dose estradiol implants did not significantly change morning corticosterone levels (One-way ANOVA, F_(2,17)_ = 0.55, *P* = 0.58; Fig 1B). Likewise, combined adrenal weight was also not different across the estradiol treatment groups (One-way ANOVA, F_(2,18)_ = 0.64, *P* = 0.54; Fig 1C). Interestingly, thymus weight, which can be influenced by both corticosterone ^44^ and estradiol ^45–47^, was significantly different between the groups (One-way ANOVA, F_(2,17)_ = 30.52, *P* < 0.0001; Fig 1D). Post-hoc Tukey’s multiple comparisons showed a significant difference between Ovx^HighE^ compared to Ovx and Ovx^LowE^ (*P* < 0.0001 and *P* = 0.0004 respectively), and a significant difference between Ovx and Ovx^LowE^ (*P* = 0.023).

**Figure 1:**
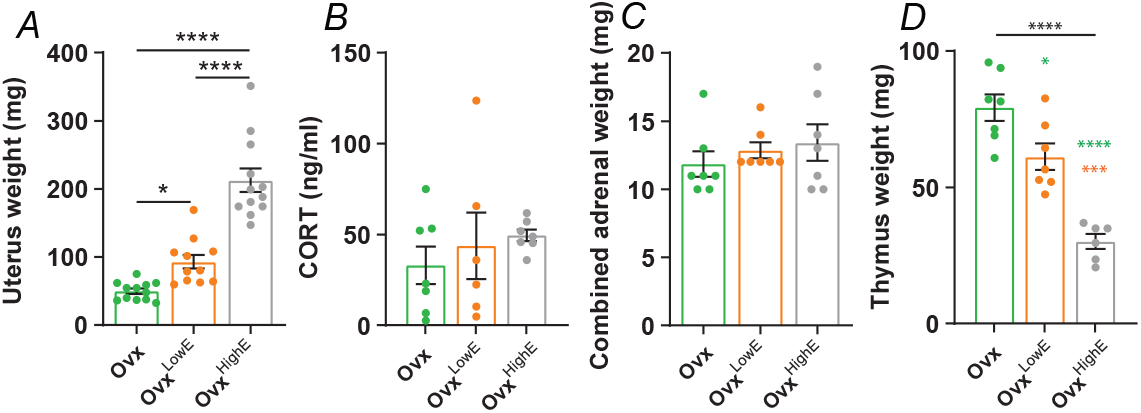
Consequences of chronic estradiol treatment. (A) Uterus weight for each group measured immediately after the animals were euthanized; Ovx (green), Ovx^LowE^ (orange) and Ovx^HighE^ (grey). Stars indicate significance identified by Tukey’s multiple comparisons, both low and high chronic estradiol implants increase uterus weight compared to Ovx animals. (B) There is no significant difference in circulating serum CORT levels taken from trunk blood samples. (C) Comparison of combined (weight of both left and right adrenal) adrenal weight between the groups. There was no significant difference in adrenal weight between the groups. (D) There is a significant difference in Thymus weight between the three groups. Black stars indicate significant result of On-way ANOVA, colored stars indicate post-hoc Tukey’s multiple comparisons test. Green stars indicate significance with Ovx group, orange stars indicate significant difference with Ovx^LowE^ group. N > 6 mice for all groups. *P* values: * ≤ 0.05, ** ≤ 0.01, *** ≤ 0.001, **** ≤ 0.0001.

### Estradiol regulates I_A_ potassium channel currents in CRH neurons

Neuronal intrinsic excitability is dictated in part by voltage-gated ion channel density and function. We have previously shown that I_A_, a transient K^+^ current, is regulated over the estrous cycle in CRH neurons ^28^. To investigate the link between estradiol levels and I_A_ currents, we used a voltage clamp protocol on CRH neurons from Ovx, Ovx^LowE^ or Ovx^HighE^ manipulated mice. Electrophysiological recordings were performed 2-3 weeks post ovariectomy. A Two-way RM ANOVA revealed that there was a significant effect of estradiol treatment on I_A_ current density (F_(2, 18)_ = 5.32, *P* = 0.015; Fig 2 A & B), a significant effect of voltage step (F_(14, 252)_ = 134.1, *P* < 0.0001) and a significant interaction (F_(28, 252)_ = 4.68, *P* < 0.0001). Post hoc tests showed that current densities at multiple voltage steps were smallest in Ovx animals compared to Ovx^LowE^ (*P* < 0.05) and Ovx^HighE^ animals (*P* < 0.05). Peak amplitude of the current at the maximum voltage step (+30 mV) was also significantly different between groups (One-way ANOVA, F_(2, 17)_ = 4.55, *P* = 0.026; Fig 2C). Post hoc comparison revealed a significant difference between Ovx and Ovx^HighE^ (*P* = 0.021) but not with Ovx^LowE^ (*P* = 0.18). These findings show that chronic estradiol manipulations lead to changes in I_A_ K^+^ currents in CRH neurons.

**Figure 2:**
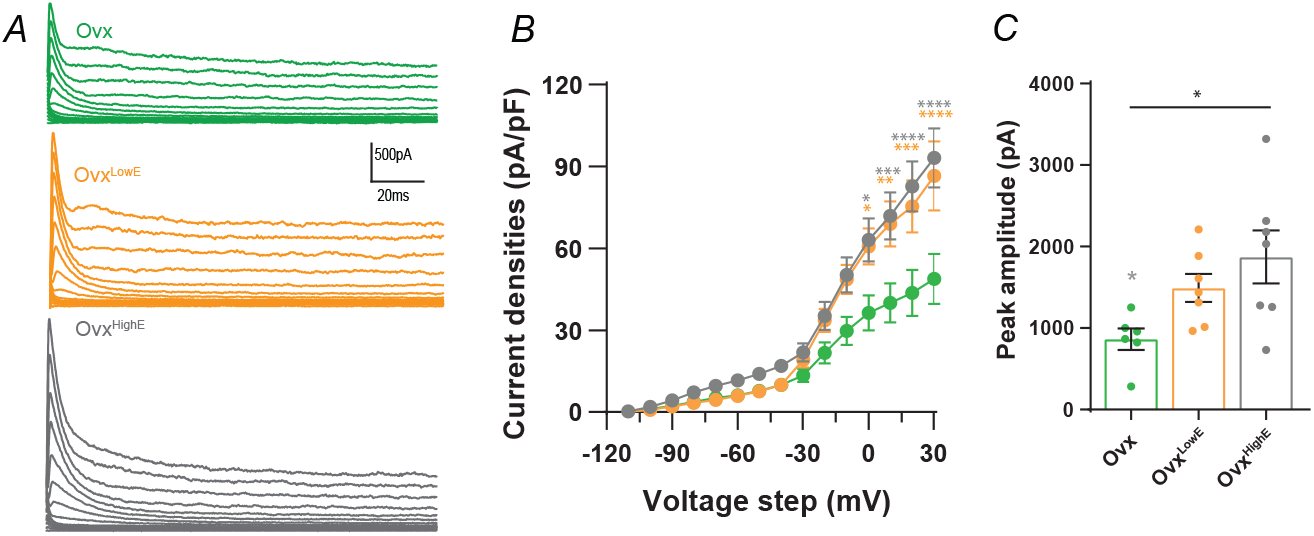
I_A_ currents are influenced by chronic estradiol treatment. (A) Evoked I_A_ currents from individual cells in each group; Ovx (green), Ovx^LowE^ (orange) and Ovx^HighE^ (grey). (B) I_A_ current densities plotted for each 10 mV voltage step from −100 to +30 mV. Cells from Ovx animals had significantly smaller I_A_ current densities compared to cells from Ovx^LowE^ and Ovx^HighE^. (C) Peak amplitude I_A_ currents (not normalized to capacitance) evoked by a +30mV step. Peak I_A_ currents in Ovx animals were significantly smaller than in Ovx^LowE^ and Ovx^HighE^. Results of one and two-way ANOVAs reported in Table 2. Star symbols denote significance by Tukey’s multiple comparisons test; orange: Ovx^LowE^ versus Ovx, grey: Ovx^HighE^ versus Ovx. There were no significant differences between Ovx^LowE^ and Ovx^HighE^ groups. N = 3 - 5 mice for all groups. *P* values: * ≤ 0.05, ** ≤ 0.01, *** ≤ 0.001, **** ≤ 0.0001.

### Estradiol regulates I_M_ potassium channel currents in CRH neurons

In addition to I_A_, M type (I_M_) potassium currents were also investigated. I_M_ currents are slowly activating, non-inactivating voltage-gated currents. They can contribute to intrinsic excitability via regulation of resting membrane potential and are ubiquitously found in neurons ^48^. A voltage clamp protocol was used to measure the relaxation of the I_M_ current (see methods). Comparison of the three groups using a Two-way RM ANOVA revealed a significant effect of chronic estradiol treatment on I_M_ current densities (F_(2,19)_ = 5.37, *P* = 0.014; Fig 3A & B). Post-hoc multiple comparisons revealed that the Ovx^HighE^ group had a significantly higher I_M_ current density compared to both Ovx (*P* = 0.04) and Ovx^LowE^ (*P* = 0.04) at the highest voltage step (−30mV). A one-way ANOVA comparing peak I_M_ current amplitude in the three groups was also significant (F_(2,19)_ = 5.01, *P* = 0.018; Fig 3C), with multiple comparisons revealing significant differences between Ovx^HighE^ and Ovx^LowE^ (*P* = 0.026) but not between Ovx^HighE^ and Ovx (*P* = 0.075). These results show that estradiol levels also regulate I_M_ K^+^channel currents in CRH neurons.

**Figure 3:**
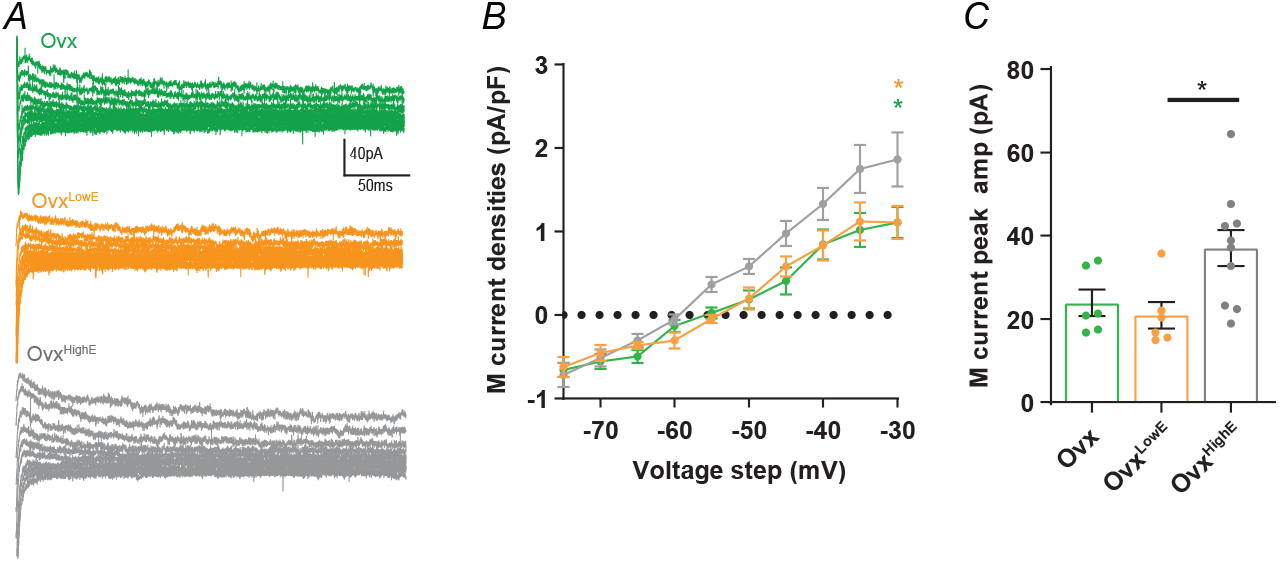
I_M_ currents are influenced by high chronic estradiol levels but not low. (A) Evoked I_M_ currents from individual cells in each group; Ovx (green), Ovx^LowE^ (orange) and Ovx^HighE^ (grey). (B) I_M_ current densities plotted for each 5 mV voltage step from −75 to −30 mV. Cells from Ovx^HighE^ animals had significantly larger I_M_ current densities compared to cells from Ovx^LowE^ and Ovx. (C) Peak amplitude I_M_ currents (not normalized to capacitance) evoked by a −30mV step. Peak I_M_ currents in Ovx^HighE^ animals were significantly larger than in Ovx^LowE^, but not compared to cells from Ovx animals. Results of one and two-way ANOVAs reported in Table 2. Star symbols denote significance by Tukey’s multiple comparisons test; orange: Ovx^LowE^ versus Ovx^HighE^, green: Ovx versus Ovx^HighE^. There were no significant differences between Ovx^LowE^ and Ovx groups. N = 3 - 5 mice for all groups. *P* values: * ≤ 0.05, ** ≤ 0.01, *** ≤ 0.001, **** ≤ 0.0001.

### Chronic estradiol manipulations do not alter CRH neuron intrinsic excitability

As both I_A_ and I_M_ currents are altered by artificially induced estradiol concentrations, we next investigated if CRH neuron intrinsic excitability was also influenced. Given the elevated K^+^ currents in CRH neurons from Ovx^LowE^ and Ovx^HighE^ mice we expected lower intrinsic excitability levels from these neurons compared to those from Ovx animals. Neurons were held around −60 mV in current clamp before injecting a family of current steps from 0 pA to +50 pA in 5 pA increments (Fig 4A and B). This protocol was used to generate a frequency/current curve (F/I curve) and was performed on CRH neurons from Ovx, Ovx^LowE^ and Ovx^HighE^ animals. CRH neuron firing frequency was not different between the groups (Two-way RM ANOVA, F_(2, 32)_ = 0.386, *P* = 0.61; Fig 4A and B, Table 1), nor was peak firing frequency (One-way ANOVA, F_(2,27)_ = 3.102, *P* = 0.68; Fig 4B insert), or the slope of the FI curves (One-way ANOVA, F_(2,32)_ = 0.19, *P* = 0.83; Table 1). First spike latency (FSL), measured from the 10pA current step onwards, was not affected by chronic estradiol treatment (Two-way RM ANOVA, F_(2,31)_ = 0.059, *P* = 0.94; Fig 4 C, Table 1). The total number of APs fired over all current steps was also similar between groups (One-way ANOVA, F_(2,30)_ = 0.17, *P* = 0.84; Fig 4D). Analysis of AP parameters showed no significant differences in amplitude, rise time, half width or decay time between the three groups (Fig 4E-H, see Table 1 for mean values and Table 2 for statistics). There were also no significant differences in capacitance or input resistance between groups (one-way ANOVA, F_(2,85)_ = 1.93, *P* = 0.15 and F_(2,85)_ = 1.6, *P* = 0.21 respectively, Table 1 and 2).

**Figure 4:**
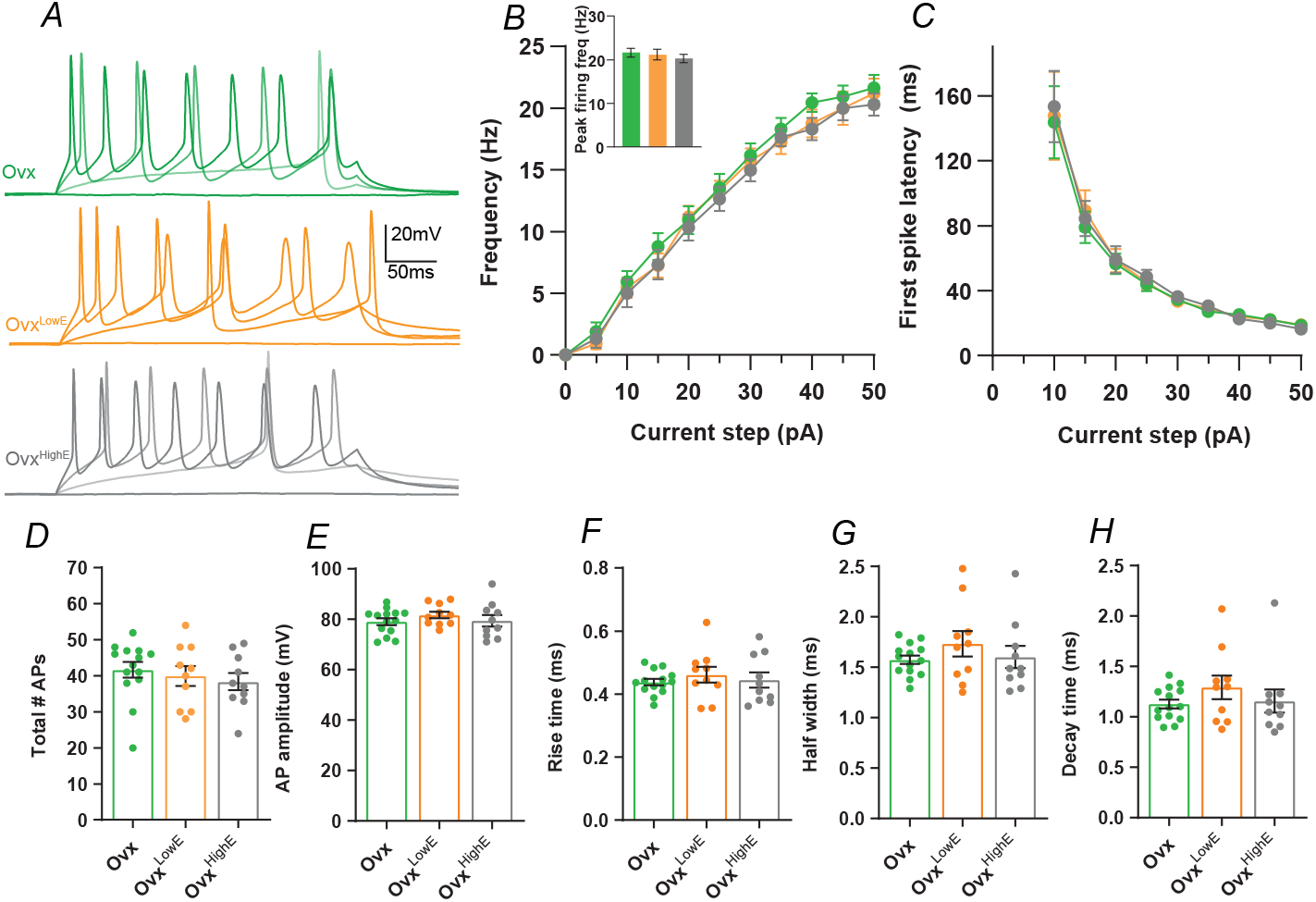
Chronic estradiol does not change CRH neuron intrinsic excitability or action potential parameters. (A) Representative responses of CRH neurons to 0 pA, 10 pA, 30 pA and 50 pA current steps of the FI curve. Ovx in green, Ovx^LowE^ in orange and Ovx^HighE^ in grey. (B) Summary data for the F/I curve. There is no significant difference between the three groups across the FI curve or at peak firing frequency (50 pA step, bar graph insert). Two-way ANOVA results reported in Table 2. (C) Graph of first spike latency (FSL) for each current step. There is no significant difference between the groups, Two-way ANOVA results reported in Table 2. (D) Bar graph of total number of APs fired over all current steps for each group. No significance between the groups. All AP parameters were measured from the first AP fired from each cell. There was no significant difference between the groups in AP amplitude (E), Rise time (F), Half width (G), or decay time (H). One-way ANOVA results reported in Table 2.

**Table 1:**

| Name | Group | Mean value ± SEM |
| --- | --- | --- |
| Total # APs fired during F/I curve | OvX | 41.64 ± 2.2 |
| | OvX<sup>LowE</sup> | 39.36 ± 2.58 |
| | OvX<sup>HighE</sup> | 42.3 ± 1.56 |
| Slope of F/I curve | OvX | 0.46 ± 0.022 Hz/pA |
| | OvX<sup>LowE</sup> | 0.45 ± 0.024 Hz/pA |
| | OvX<sup>HighE</sup> | 0.44 ± 0.019 Hz/pA |
| Peak firing rate (50pA step) | OvX | 21.65 ± 1.03 Hz |
| | OvX<sup>LowE</sup> | 21.09 ± 1.21 Hz |
| | OvX<sup>HighE</sup> | 20.31 ± 0.92 Hz |
| AP amplitude | OvX | 79.11 ± 1.35 pA |
| | OvX<sup>LowE</sup> | 81.77 ± 1.35 pA |
| | OvX<sup>HighE</sup> | 79.03 ± 2.09 pA |
| AP rise time | OvX | 0.44 ± 0.01 ms |
| | OvX<sup>LowE</sup> | 0.46 ± 0.03 ms |
| | OvX<sup>HighE</sup> | 0.45 ± 0.22 ms |
| AP half width | OvX | 1.58 ± 0.04 ms |
| | OvX<sup>LowE</sup> | 1.74 ± 0.13 ms |
| | OvX<sup>HighE</sup> | 1.6 ± 0.1 ms |
| AP decay time | OvX | 1.13 ± 0.04 ms |
| | OvX<sup>LowE</sup> | 1.29 ± 0.18 ms |
| | OvX<sup>HighE</sup> | 1.14 ± 0.11 ms |
| Input resistance | OvX | 1.7 ± 0.11 GΩ |
| | OvX<sup>LowE</sup> | 1.99 ± 0.14 GΩ |
| | OvX<sup>HighE</sup> | 1.76 ± 0.11 GΩ |
| Capacitance | OvX | 21.06 ± 1.13 pF |
| | OvX<sup>LowE</sup> | 18.49 ± 0.99 pF |
| | OvX<sup>HighE</sup> | 20.63 ± 0.78 pF |
| Uterus weight | OvX | 50 ± 3.84 mg (*) (****) |
| | OvX<sup>LowE</sup> | 93.18 ± 10.16 mg (****) |
|  | Ovx<sup>HighE</sup> | 212.58 ± 17.18 mg |
| Adrenal weight | Ovx | 11.85 ± 0.94 mg |
|  | Ovx<sup>LowE</sup> | 12.86 ± 0.59 mg |
|  | Ovx<sup>HighE</sup> | 13.43 ± 1.32 mg |
| Serum CORT concentration | Ovx | 33.08 ± 10.31 ng/ml |
|  | Ovx<sup>LowE</sup> | 43.76 ± 18.3 ng/ml |
|  | Ovx<sup>HighE</sup> | 49.55 ± 3.22 ng/ml |
| Thymus weight | Ovx | 79.27 ± 4.88 mg (*) (****) |
|  | Ovx<sup>LowE</sup> | 61.26 ± 4.8 mg (***) |
|  | Ovx<sup>HighE</sup> | 30.13 ± 2.75 mg |

**Table 2:**
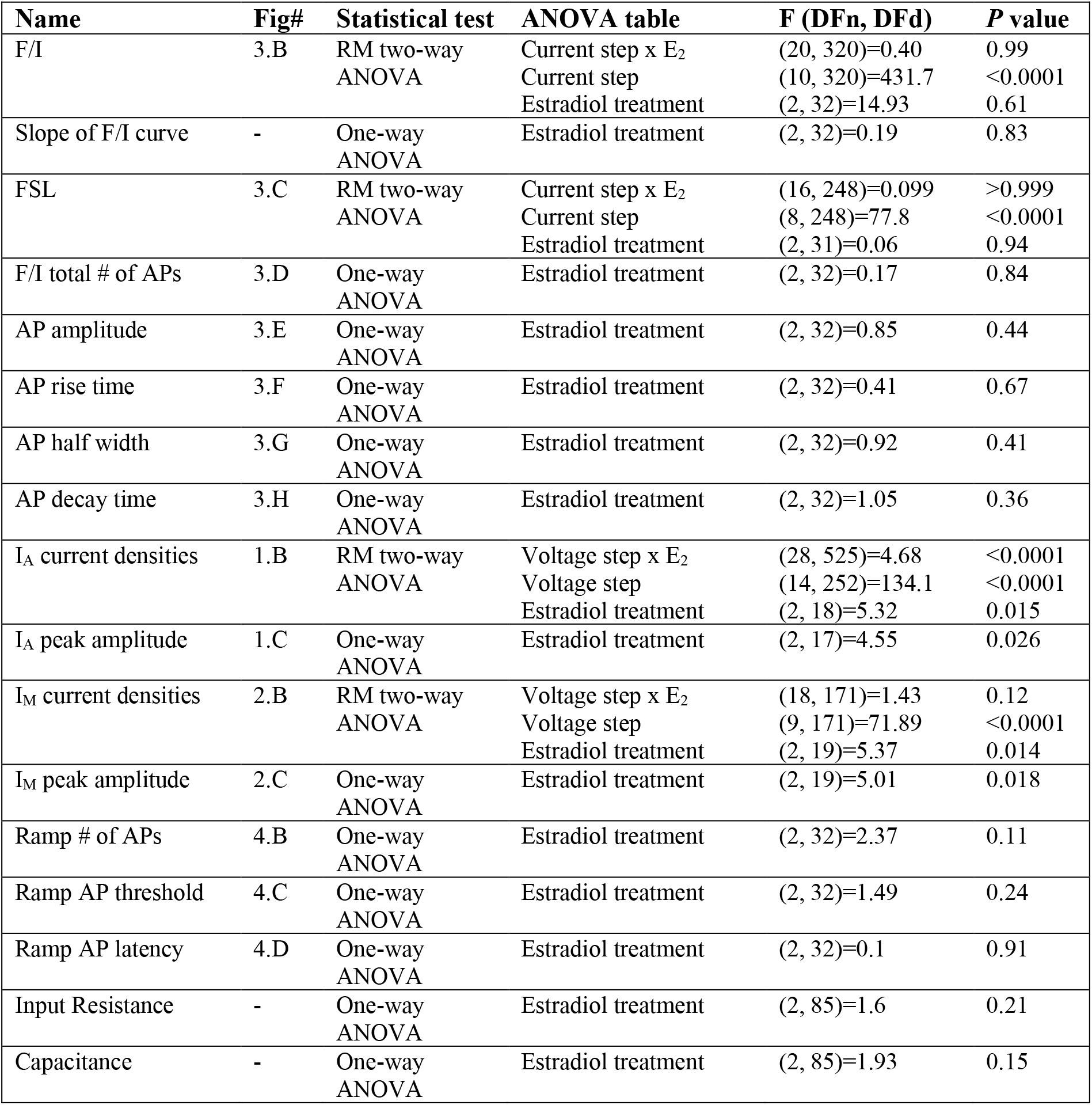
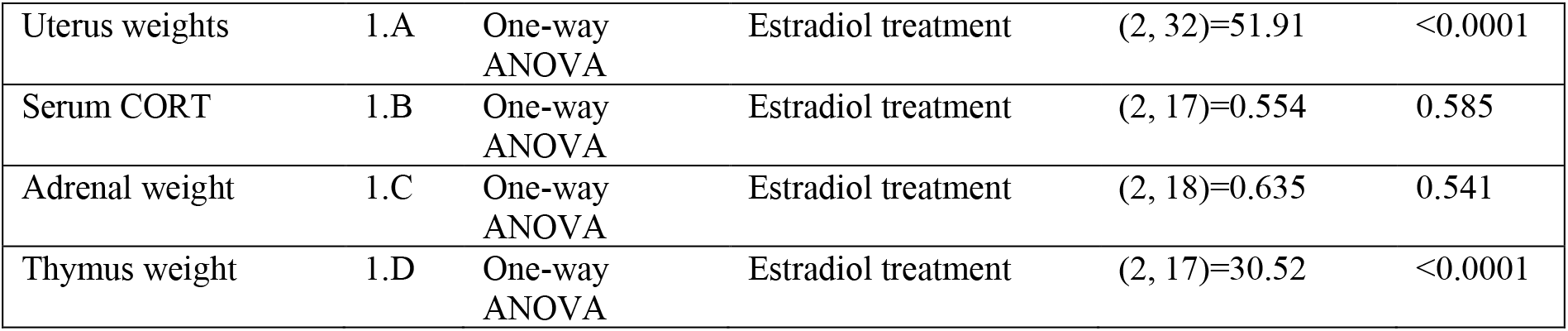

| Name | Fig# | Statistical test | ANOVA table | F (DFn, DFd) | P value |
| --- | --- | --- | --- | --- | --- |
| F/I | 3.B | RM two-way ANOVA | Current step x E<sub>2</sub> | (20, 320)=0.40 | 0.99 |
|  |  |  | Current step | (10, 320)=431.7 | <0.0001 |
|  |  |  | Estradiol treatment | (2, 32)=14.93 | 0.61 |
| Slope of F/I curve | - | One-way ANOVA | Estradiol treatment | (2, 32)=0.19 | 0.83 |
| FSL | 3.C | RM two-way ANOVA | Current step x E<sub>2</sub> | (16, 248)=0.099 | >0.999 |
|  |  |  | Current step | (8, 248)=77.8 | <0.0001 |
|  |  |  | Estradiol treatment | (2, 31)=0.06 | 0.94 |
| F/I total # of APs | 3.D | One-way ANOVA | Estradiol treatment | (2, 32)=0.17 | 0.84 |
| AP amplitude | 3.E | One-way ANOVA | Estradiol treatment | (2, 32)=0.85 | 0.44 |
| AP rise time | 3.F | One-way ANOVA | Estradiol treatment | (2, 32)=0.41 | 0.67 |
| AP half width | 3.G | One-way ANOVA | Estradiol treatment | (2, 32)=0.92 | 0.41 |
| AP decay time | 3.H | One-way ANOVA | Estradiol treatment | (2, 32)=1.05 | 0.36 |
| I<sub>A</sub> current densities | 1.B | RM two-way ANOVA | Voltage step x E<sub>2</sub> | (28, 525)=4.68 | <0.0001 |
|  |  |  | Voltage step | (14, 252)=134.1 | <0.0001 |
|  |  |  | Estradiol treatment | (2, 18)=5.32 | 0.015 |
| I<sub>A</sub> peak amplitude | 1.C | One-way ANOVA | Estradiol treatment | (2, 17)=4.55 | 0.026 |
| I<sub>M</sub> current densities | 2.B | RM two-way ANOVA | Voltage step x E<sub>2</sub> | (18, 171)=1.43 | 0.12 |
|  |  |  | Voltage step | (9, 171)=71.89 | <0.0001 |
|  |  |  | Estradiol treatment | (2, 19)=5.37 | 0.014 |
| I<sub>M</sub> peak amplitude | 2.C | One-way ANOVA | Estradiol treatment | (2, 19)=5.01 | 0.018 |
| Ramp # of APs | 4.B | One-way ANOVA | Estradiol treatment | (2, 32)=2.37 | 0.11 |
| Ramp AP threshold | 4.C | One-way ANOVA | Estradiol treatment | (2, 32)=1.49 | 0.24 |
| Ramp AP latency | 4.D | One-way ANOVA | Estradiol treatment | (2, 32)=0.1 | 0.91 |
| Input Resistance | - | One-way ANOVA | Estradiol treatment | (2, 85)=1.6 | 0.21 |
| Capacitance | - | One-way ANOVA | Estradiol treatment | (2, 85)=1.93 | 0.15 |
| Uterus weights | 1.A | One-way<br>ANOVA | Estradiol treatment | (2, 32)=51.91 | <0.0001 |
| Serum CORT | 1.B | One-way<br>ANOVA | Estradiol treatment | (2, 17)=0.554 | 0.585 |
| Adrenal weight | 1.C | One-way<br>ANOVA | Estradiol treatment | (2, 18)=0.635 | 0.541 |
| Thymus weight | 1.D | One-way<br>ANOVA | Estradiol treatment | (2, 17)=30.52 | <0.0001 |

In addition to an F/I curve, CRH neuron excitability was also tested using a current clamp ramp protocol consisting of a 40pA ramp delivered over 1 second (Figure 5A). This protocol gives a more accurate measurement of latency to first spike and AP threshold compared to measurements from F/I curves. Neither the number of APs fired (Fig 5B), AP threshold (Fig 5C) or first spike latency (Fig 5D) were significantly different between the groups (One-way ANOVA, F_(2,29)_ = 1.89, *P* = 0.17, F_(2,31)_ = 0.64, *P* = 0.54, F_(2,31)_ = 0.22, *P* = 0.80, respectively). Despite chronic estradiol treatment causing changes in K^+^ channel function, these results show that chronic estradiol manipulation had no impact on CRH neuron intrinsic excitability.

**Figure 5:**
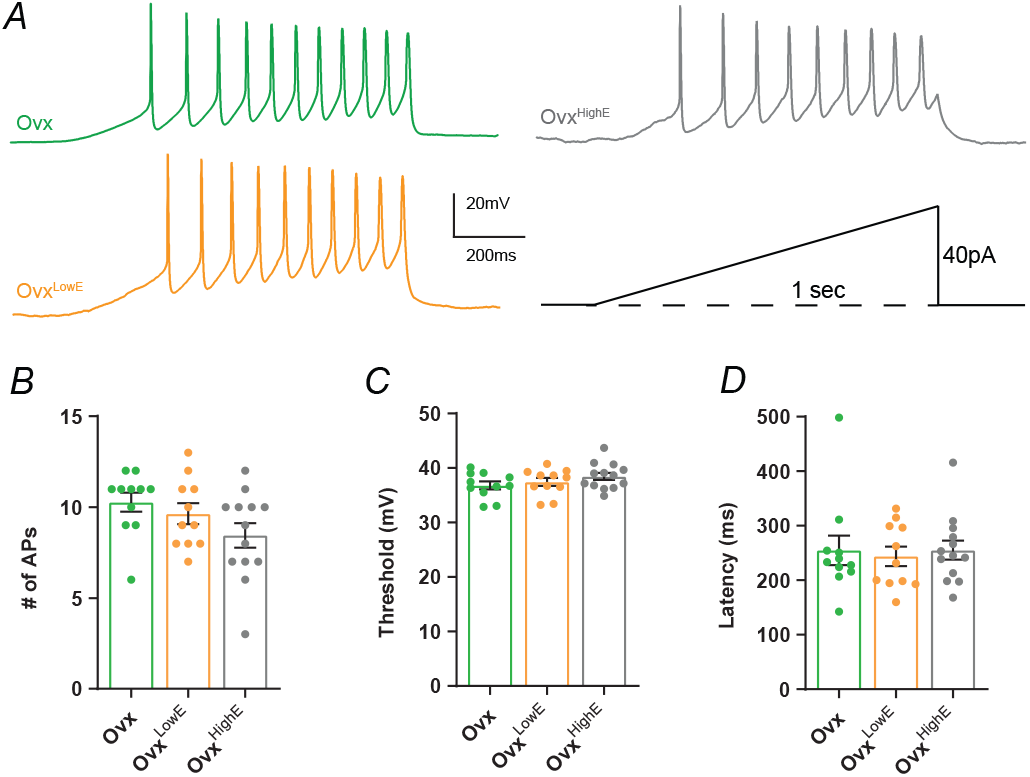
Chronic estradiol does not influence CRH neuron intrinsic excitability measured by a current ramp. (A) Example traces showing CRH neuron spiking response to a one second, 40 pA, current ramp protocol (bottom right). Green is Ovx, orange is Ovx^LowE^, grey is Ovx^HighE^. (B) No significant difference in the total number of APs fired during the ramp protocol between the three groups. (C) Neither high nor low estradiol concentrations alter AP threshold compared to Ovx. AP threshold was defined as the voltage at which the AP first derivative crossed 10 mV/ms. (D) There is no change in AP latency between the groups. Results of One-way ANOVAs are reported in Table 2.

### I_A_ currents correlate with CRH neuron excitability

We have previously shown that I_A_ currents are regulated over the female estrous cycle and control CRH neuron intrinsic excitability ^28^. We took the data from this previous work along with data from this current study and performed Pearson’s correlation tests for I_A_ current density versus various parameters of excitability measured from F/I curves and Ramp protocol (Figure 6). We included data from the following groups: intact estrus, intact proestrus, intact diestrus, Ovx, Ovx^LowE^ and Ovx^HighE^.

**Figure 6:**
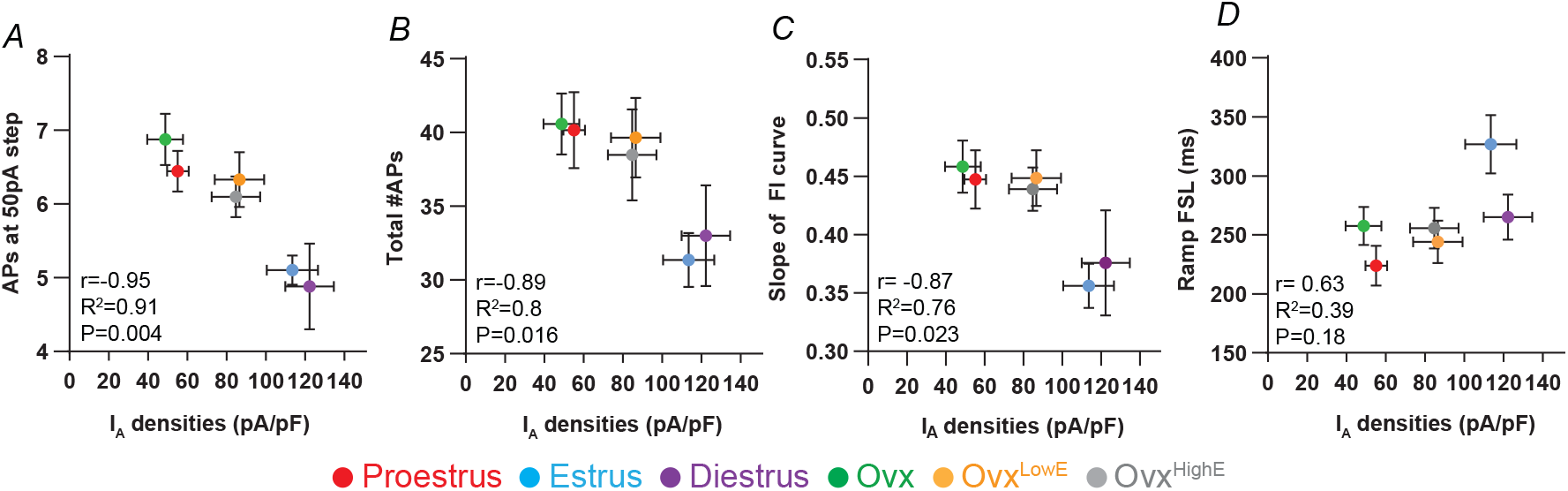
I_A_ current densities correlate strongly with measures of CRH neuron excitability. (A) Correlation analysis between the number of APs fired at the highest (50 pA) current step given during the FI curve and peak I_A_ current densities. Red is proestrus, blue is estrus, purple is diestrus, green is Ovx, orange is Ovx^LowE^ and grey is Ovx^HighE^ (B) I_A_ current densities versus total number of APs fired over the entire FI curve for each group. (C) The slope of FI curve correlated to I_A_ current densities. (D) First spike latency (FSL) of current Ramp protocol correlated to I_A_ current densities. R, R^2^ and P values for each comparison are listed on individual graphs.

I_A_ current densities were found to be negatively correlated with the number of APs fired at the 50 pA current step (r = 0.95, *P* = 0.002; Fig 6A), the total number of APs fired (r = 0.89, *P* = 0.016; Fig 6B), and the slope of the FI curve (r = 0.87, *P* = 0.023; Fig 6C). I_A_ current densities were also positively correlated with current Ramp FSL, although this was not significant (r = 0.63, *P* = 0.18; Fig 6D). These data show that changes in I_A_ current density in CRH neurons are correlated with several parameters of intrinsic excitability.

## Discussion

Circulating levels of estradiol have been previously shown to regulate the HPA axis ^14,49,50^, however, the impact of estradiol on CRH neuron excitability has been less clear. In the current study, we find that compared to Ovx mice, replacement with either low or high doses of estradiol increased I_A_ current density in CRH neurons. For I_M_ currents, only high estradiol concentrations led to an increase in current density. Despite these changes in K^+^ currents following estradiol manipulations, there were no significant changes in intrinsic excitability parameters. However, when we combined data from this current study with that of previous work looking at excitability in CRH neurons from intact, cycling females, we found significant correlations between I_A_ current density and several measures of excitability.

These findings differ compared with previous studies investigating estradiol effects on K^+^ currents in hypothalamic neurons. Hu et al. demonstrated that acute bath application of estradiol (100 nM, 10 minutes) onto CRH neurons from Ovx mice could supresses I_M_ currents ^39^. This effect could be replicated with a membrane-associated estrogen receptor (ER) agonist suggesting the fast suppression of M currents by estradiol was mediated via a non-genomic signalling mechanism ^39^. In other neural populations, chronic estradiol replacement in Ovx animals has also been shown to suppress K^+^ currents. In rostral ventrolateral medulla projecting preautonomic PVN neurons, estradiol treatment in Ovx rats was sufficient to reduce I_A_ current density ^51^. In GnRH neurons, estradiol treatment in Ovx mice also decreased both I_A_ and IK currents ^31^. Previously, we have shown that I_A_ currents in CRH neurons are smallest during the proestrus phase and largest during the estrus stage of the mouse estrous cycle ^28^. However, hormone profiles in intact animals will be different compared to Ovx animals with estradiol replacement and this may underly the differing findings.

What signalling pathways could be responsible for the effects of estradiol on K^+^ channel function in CRH neurons in this current study? Estradiol acts through two main receptors, ERα and ERβ. In Ovx rats these receptors have opposing effects on stress induced glucocorticoid secretion with ERα increasing secretion while ERβ decreases it ^52,53^. ERβ is expressed in PVN neurons and shows colocalization with CRH ^54,55^. Therefore, estradiol acting through ERβ could possibly be mediating the effects observed. Comparatively, ERα shows little to no expression in mouse PVN CRH neurons ^56^, however, may regulate CRH neuron function indirectly via afferent inputs ^17^. There are a number of different neural populations that express ERα and project to the PVN including neurons in the arcuate nucleus ^57,58^, the bed nucleus of stria terminalis ^59^ and the peri-PVN region ^52^. In addition, estradiol manipulations are known to regulate the signalling of other neurotransmitter systems in the PVN including serotonin ^60^, oxytocin ^61^ and vasopressin ^62,63^. In summary, because estradiol modulates a number neural circuits, neurotransmitter systems and hormone systems, it is likely that changes in CRH neuron function result from a combination of direct and indirect effects of estradiol. While it is currently unclear the relative importance of each of these pathways for mediating the changes in K^+^ channel function in CRH neurons, we can conclude that the initial trigger for these changes is the change in circulating estradiol.

Despite changes in K^+^ channel activity, chronic estradiol manipulations did not influence specific parameters of CRH neuron intrinsic excitability. However, the correlation analysis showed that there is a significant relationship between the I_A_ currents and CRH excitability when data from intact, cycling animals was included. Interestingly, while estradiol replacement increases I_A_ current density in Ovx^LowE^ and Ovx^HighE^ animals compared to Ovx, I_A_ current density does not reach the same level as that seen in intact diestrus or estrus mice. This data suggests that the magnitude of increase in K^+^ current density following estradiol replacement may not have been large enough to change CRH neuron intrinsic excitability.

A second reason why estradiol may not have impacted CRH neuron intrinsic excitability is homeostatic plasticity. Past research has shown that chronic manipulations of K^+^ channel function can induce compensatory changes in excitability known as homeostatic plasticity ^64^. This form of plasticity acts to return the activity of neural circuits to a homeostatic set point ^64–68^. CRH neurons may similarly have an “homeostatic setpoint” firing rate, which in the intact animal is subject to a dynamically changing hormonal environment resulting in temporary changes in excitability across the estrous cycle ^28^. However, in a static hormonal environment, like that seen in Ovx + estradiol treated mice, CRH neuron spiking excitability may retune to the setpoint despite differences in K^+^ channel activity. In order for this to happen, the function of other ion channels would need to be regulated. This hypothesis would be interesting to address in future work.

Together, data from this current study shows that chronic estradiol elevations lead to enhanced potassium channel currents in CRH neurons. While there were no obvious effects on spiking excitability, we predict that enhanced K^+^ channel function may impact how these neurons integrate and process stress relevant synaptic inputs.

## Acknowledgements

This work was supported by a Royal Society of New Zealand Marsden Grant. The authors would also like to thank Shaojie Zheng and Dr Joon Kim for assistance with this project.

## References

1. Herman JP, Cullinan WE. Neurocircuitry of stress: central control of the hypothalamo–pituitary–adrenocortical axis. Trends in neurosciences. 1997;20(2):78–84.

2. Ulrich-Lai YM, Herman JP. Neural regulation of endocrine and autonomic stress responses. Nature reviews neuroscience. 2009;10(6):397–409.

3. Kim JS, Han SY, Iremonger KJ. Stress experience and hormone feedback tune distinct components of hypothalamic CRH neuron activity. Nature communications. 2019;10(1):1–15.

4. Füzesi T, Daviu N, Wamsteeker Cusulin JI, Bonin RP, Bains JS. Hypothalamic CRH neurons orchestrate complex behaviours after stress. Nature communications. 2016;7:11937.

5. Kim J, Lee S, Fang Y-Y, et al. Rapid, biphasic CRF neuronal responses encode positive and negative valence. Nature neuroscience. 2019;22(4):576–585.

6. Sterley T-L, Baimoukhametova D, Füzesi T, et al. Social transmission and buffering of synaptic changes after stress. Nature neuroscience. 2018;21(3):393–403.

7. Daviu N, Füzesi T, Rosenegger DG, et al. Paraventricular nucleus CRH neurons encode stress controllability and regulate defensive behavior selection. Nature Neuroscience. 2020;23(3):398–410.

8. Seale J, Wood, SA, Atkinson, HC, Bate, E, Lightman, SL, Ingram, CD, Jessop, DS, Harbuz, MS Gonadectomy reverses the sexually diergic patterns of circadian and stress-induced hypothalamic-pituitary-adrenal axis activity in male and female rats. J Neuroendocrinol. 2004;16:516–24.

9. Atkinson HC, Waddell BJ. Circadian variation in basal plasma corticosterone and adrenocorticotropin in the rat: sexual dimorphism and changes across the estrous cycle. Endocrinology. 1997;138(9):3842–3848.

10. Viau V, Meaney MJ. Variations in the hypothalamic-pituitary-adrenal response to stress during the estrous cycle in the rat. Endocrinology. 1991;129(5):2503–2511.

11. Nilsson ME, Vandenput L, Tivesten Å, et al. Measurement of a comprehensive sex steroid profile in rodent serum by high-sensitive gas chromatography-tandem mass spectrometry. Endocrinology. 2015;156(7):2492–2502.

12. Babb JA, Masini CV, Day HE, Campeau S. Sex differences in activated corticotropin-releasing factor neurons within stress-related neurocircuitry and hypothalamic–pituitary–adrenocortical axis hormones following restraint in rats. Neuroscience. 2013;234:40–52.

13. Young EA, Altemus M, Parkison V, Shastry S. Effects of estrogen antagonists and agonists on the ACTH response to restraint stress in female rats. Neuropsychopharmacology. 2001;25(6):881–891.

14. Figueiredo HF, Ulrich-Lai YM, Choi DC, Herman JP. Estrogen potentiates adrenocortical responses to stress in female rats. American Journal of Physiology-Endocrinology and Metabolism. 2007;292(4):E1173–E1182.

15. Kitay JI. Effect of oestradiol in adrenal corticoidogenesis: an additional step in steroid biosynthesis. Nature. 1966;209(5025):808–809.

16. Lo MJ, Chang LL, Wang PS. Effects of estradiol on corticosterone secretion in ovariectomized rats. Journal of cellular biochemistry. 2000;77(4):560–568.

17. Dayas C, Xu Y, Buller K, Day T. Effects of chronic oestrogen replacement on stress-induced activation of hypothalamic-pituitary-adrenal axis control pathways. Journal of neuroendocrinology. 2000;12(8):784–794.

18. Gerrits M, Grootkarijn A, Bekkering BF, Bruinsma M, Den Boer JA, Ter Horst GJ. Cyclic estradiol replacement attenuates stress-induced c-Fos expression in the PVN of ovariectomized rats. Brain research bulletin. 2005;67(1-2):147–155.

19. Kreisman MJ, McCosh RB, Tian K, Song CI, Breen KM. Estradiol enables chronic corticosterone to inhibit pulsatile luteinizing hormone secretion and suppress Kiss1 neuronal activation in female mice. Neuroendocrinology. 2020;110(6):501–516.

20. Aoki M, Shimozuru M, Kikusui T, Takeuchi Y, Mori Y. Sex differences in behavioral and corticosterone responses to mild stressors in ICR mice are altered by ovariectomy in peripubertal period. Zoological science. 2010;27(10):783–789.

21. Speert DB, McClennen SJ, Seasholtz AF. Sexually dimorphic expression of corticotropin-releasing hormone-binding protein in the mouse pituitary. Endocrinology. 2002;143(12):4730–4741.

22. Wada T, Sameshima A, Yonezawa R, et al. Impact of central and peripheral estrogen treatment on anxiety and depression phenotypes in a mouse model of postmenopausal obesity. PLoS One. 2018;13(12):e0209859.

23. Daodee S, Monthakantirat O, Tantipongpiradet A, et al. Effect of Yakae-Prajamduen-Jamod Traditional Thai Remedy on Cognitive Impairment in an Ovariectomized Mouse Model and Its Mechanism of Action. Molecules. 2022;27(13):4310.

24. Eid RS, Lieblich SE, Duarte-Guterman P, et al. Selective activation of estrogen receptors α and β: Implications for depressive-like phenotypes in female mice exposed to chronic unpredictable stress. Hormones and behavior. 2020;119:104651.

25. Tantipongpiradet A, Monthakantirat O, Vipatpakpaiboon O, et al. Effects of puerarin on the ovariectomy-induced depressive-like behavior in ICR mice and its possible mechanism of action. Molecules. 2019;24(24):4569.

26. Ghobadi N, Sahraei H, Meftahi GH, Bananej M, Salehi S. Effect of estradiol replacement in ovariectomized NMRI mice in response to acute and chronic stress. Journal of applied pharmaceutical science. 2016;6(11):176–184.

27. Tang AC, Nakazawa M, Romeo RD, Reeb BC, Sisti H, McEwen BS. Effects of long-term estrogen replacement on social investigation and social memory in ovariectomized C57BL/6 mice. Hormones and behavior. 2005;47(3):350–357.

28. Power EM, Iremonger KJ. Plasticity of intrinsic excitability across the estrous cycle in hypothalamic CRH neurons. Scientific reports. 2021;11(1):1–7.

29. Arroyo A, Kim BS, Biehl A, Yeh J, Bett GC. Expression of kv4. 3 voltage-gated potassium channels in rat gonadotrophin-releasing hormone (GnRH) neurons during the estrous cycle. Reproductive Sciences. 2011;18(2):136–144.

30. Vastagh C, Solymosi N, Farkas I, Liposits Z. Proestrus differentially regulates expression of ion channel and calcium homeostasis genes in GnRH neurons of mice. Frontiers in molecular neuroscience. 2019;12:137.

31. DeFazio RA, Moenter SM. Estradiol feedback alters potassium currents and firing properties of gonadotropin-releasing hormone neurons. Molecular endocrinology. 2002;16(10):2255–2265.

32. Taniguchi H, He M, Wu P, et al. A resource of Cre driver lines for genetic targeting of GABAergic neurons in cerebral cortex. Neuron. 2011;71(6):995–1013.

33. Chen Y, Molet J, Gunn BG, Ressler K, Baram TZ. Diversity of reporter expression patterns in transgenic mouse lines targeting corticotropin-releasing hormone-expressing neurons. Endocrinology. 2015;156(12):4769–4780.

34. Wamsteeker-Cusulin JI, Füzesi T, Watts AG, Bains JS. Characterization of corticotropin-releasing hormone neurons in the paraventricular nucleus of the hypothalamus of Crh-IRES-Cre mutant mice. PLoS One. 2013;8(5):e64943.

35. Jamieson B, Nair B, Iremonger K. Regulation of hypothalamic corticotropin-releasing hormone neurone excitability by oxytocin. Journal of neuroendocrinology. 2017;29(11):e12532.

36. Desroziers E, Brock O, Bakker J. Potential contribution of progesterone receptors to the development of sexual behavior in male and female mice. Hormones and behavior. 2017;90:31–38.

37. Hellier V, Brock O, Candlish M, et al. Female sexual behavior in mice is controlled by kisspeptin neurons. Nature communications. 2018;9(1):1–12.

38. Porteous R, Haden P, Hackwell EC, et al. Reformulation of PULSAR for Analysis of Pulsatile LH Secretion and a Revised Model of Estrogen-Negative Feedback in Mice. Endocrinology. 2021;162(11):bqab165.

39. Hu P, Liu J, Yasrebi A, et al. Gq protein-coupled membrane-initiated estrogen signaling rapidly excites corticotropin-releasing hormone neurons in the hypothalamic paraventricular nucleus in female mice. Endocrinology. 2016;157(9):3604–3620.

40. Roepke TA, Qiu J, Smith AW, Rønnekleiv OK, Kelly MJ. Fasting and 17β-estradiol differentially modulate the M-current in neuropeptide Y neurons. Journal of Neuroscience. 2011;31(33):11825–11835.

41. Owens JW, Ashby J. Critical review and evaluation of the uterotrophic bioassay for the identification of possible estrogen agonists and antagonists: in support of the validation of the OECD uterotrophic protocols for the laboratory rodent. Critical reviews in toxicology. 2002;32(6):445–520.

42. Serova LI, Harris HA, Maharjan S, Sabban EL. Modulation of several responses to stress by estradiol benzoate and selective estrogen receptor agonists. The Journal of endocrinology. 2010;205(3):253.

43. Kitay JI. Effects of estradiol on pituitary-adrenal function in male and female rats. Endocrinology. 1963;72(6):947–954.

44. Karatsoreos IN, Bhagat SM, Bowles NP, Weil ZM, Pfaff DW, McEwen BS. Endocrine and physiological changes in response to chronic corticosterone: a potential model of the metabolic syndrome in mouse. Endocrinology. 2010;151(5):2117–2127.

45. Clarke AG, Kendall MD. Histological changes in the thymus during mouse pregnancy. Thymus. 1989;14(1-3):65–78.

46. Utsuyama M, Hirokawa K. Hypertrophy of the thymus and restoration of immune functions in mice and rats by gonadectomy. Mechanisms of ageing and development. 1989;47(3):175–185.

47. Zoller AL, Kersh GJ. Estrogen induces thymic atrophy by eliminating early thymic progenitors and inhibiting proliferation of β-selected thymocytes. The Journal of Immunology. 2006;176(12):7371–7378.

48. Gutman GA, Chandy KG, Grissmer S, et al. International Union of Pharmacology. LIII. Nomenclature and molecular relationships of voltage-gated potassium channels. Pharmacological reviews. 2005;57(4):473–508.

49. Patchev VK, Hayashi S, Orikasa C, Almeida OF. Implications of estrogen-dependent brain organization for gender differences in hypothalamo-pituitary-adrenal regulation. The FASEB journal. 1995;9(5):419–423.

50. Roy BN, Reid RL, Van Vugt DA. The effects of estrogen and progesterone on corticotropin-releasing hormone and arginine vasopressin messenger ribonucleic acid levels in the paraventricular nucleus and supraoptic nucleus of the rhesus monkey. Endocrinology. 1999;140(5):2191–2198.

51. Lee SK, Ryu PD, Lee SY. Estrogen replacement modulates voltage-gated potassium channels in rat presympathetic paraventricular nucleus neurons. BMC neuroscience. 2013;14(1):134.

52. Weiser M, Handa RJ. Estrogen impairs glucocorticoid dependent negative feedback on the hypothalamic–pituitary–adrenal axis via estrogen receptor alpha within the hypothalamus. Neuroscience. 2009;159(2):883–895.

53. Weiser MJ, Foradori CD, Handa RJ. Estrogen receptor beta activation prevents glucocorticoid receptor-dependent effects of the central nucleus of the amygdala on behavior and neuroendocrine function. Brain research. 2010;1336:78–88.

54. Lund TD, Hinds LR, Handa RJ. The androgen 5α-dihydrotestosterone and its metabolite 5α-androstan-3β, 17β-diol inhibit the hypothalamo–pituitary–adrenal response to stress by acting through estrogen receptor β-expressing neurons in the hypothalamus. Journal of Neuroscience. 2006;26(5):1448–1456.

55. Oyola MG, Thompson MK, Handa AZ, Handa RJ. Distribution and chemical composition of estrogen receptor β neurons in the paraventricular nucleus of the female and male mouse hypothalamus. Journal of Comparative Neurology. 2017;525(17):3666–3682.

56. Suzuki S, Handa RJ. Estrogen receptor-β, but not estrogen receptor-α, is expressed in prolactin neurons of the female rat paraventricular and supraoptic nuclei: Comparison with other neuropeptides. Journal of Comparative Neurology. 2005;484(1):28–42.

57. Franceschini I, Lomet D, Cateau M, Delsol G, Tillet Y, Caraty A. Kisspeptin immunoreactive cells of the ovine preoptic area and arcuate nucleus co-express estrogen receptor alpha. Neuroscience letters. 2006;401(3):225–230.

58. Handa RJ, Weiser MJ. Gonadal steroid hormones and the hypothalamo–pituitary–adrenal axis. Frontiers in neuroendocrinology. 2014;35(2):197–220.

59. Shughrue PJ, Lane MV, Merchenthaler I. Comparative distribution of estrogen receptor-α and-β mRNA in the rat central nervous system. Journal of Comparative Neurology. 1997;388(4):507–525.

60. McAllister CE, Creech R, Kimball P, Muma NA, Li Q. GPR30 is necessary for estradiol-induced desensitization of 5-HT1A receptor signaling in the paraventricular nucleus of the rat hypothalamus. Psychoneuroendocrinology. 2012;37(8):1248–1260.

61. Amico JA, Seif SM, Robinson AG. Oxytocin in human plasma: correlation with neurophysin and stimulation with estrogen. The Journal of Clinical Endocrinology & Metabolism. 1981;52(5):988–993.

62. Lagunas N, Marraudino M, de Amorim M, et al. Estrogen receptor beta and G protein-coupled estrogen receptor 1 are involved in the acute estrogenic regulation of arginine-vasopressin immunoreactive levels in the supraoptic and paraventricular hypothalamic nuclei of female rats. Brain Research. 2019;1712:93–100.

63. Vilhena-Franco T, Mecawi AS, Almeida-Pereira G, Lucio-Oliveira F, Elias LLK, Antunes-Rodrigues J. Oestradiol acts through its beta receptor to increase vasopressin neuronal activation and secretion induced by dehydration. Journal of Neuroendocrinology. 2019;31(4):e12712.

64. Burrone J, O’Byrne M, Murthy VN. Multiple forms of synaptic plasticity triggered by selective suppression of activity in individual neurons. Nature. 2002;420(6914):414–418.

65. Hengen KB, Lambo ME, Van Hooser SD, Katz DB, Turrigiano GG. Firing rate homeostasis in visual cortex of freely behaving rodents. Neuron. 2013;80(2):335–342.

66. Hengen KB, Pacheco AT, McGregor JN, Van Hooser SD, Turrigiano GG. Neuronal firing rate homeostasis is inhibited by sleep and promoted by wake. Cell. 2016;165(1):180–191.

67. Keck T, Keller GB, Jacobsen RI, Eysel UT, Bonhoeffer T, Hübener M. Synaptic scaling and homeostatic plasticity in the mouse visual cortex in vivo. Neuron. 2013;80(2):327–334.

68. Turrigiano GG, Leslie KR, Desai NS, Rutherford LC, Nelson SB. Activity-dependent scaling of quantal amplitude in neocortical neurons. Nature. 1998;391(6670):892–896.

